# Discovery of novel antimicrobial resistance genes in food and fertiliser using a high-throughput gene capture and functional screening platform

**DOI:** 10.64898/2026.03.15.711940

**Authors:** Vaheesan Rajabal, Timothy M. Ghaly, Elena Colombi, Dylan H. Russell, Caleb Sia, Bhumika Shah, Vanessa J. McPherson, Qin Qi, Nicholas V. Coleman, Michael R. Gillings, Sasha G. Tetu

**Affiliations:** School of Natural Sciences, Macquarie University, New South Wales, 2109; ARC Centre of Excellence in Synthetic Biology, Macquarie University, New South Wales, 2109

**Keywords:** Antimicrobial resistance, One Health, Integron, Mobile genetic elements, Functional metagenomics, Environmental resistome, Environmental surveillance

## Abstract

Integrons are genetic elements that drive bacterial adaptation by capturing and expressing mobile gene cassettes. They play a key role in dissemination of antimicrobial resistance (AMR) genes, particularly in Gram-negative bacteria. In addition to known AMR determinants, integron gene cassettes carry a vast reservoir of novel genes whose functions are largely uncharacterised, making it difficult to assess their full contribution to the resistome. Contributing to this are limitations in current sequence-based prediction methods which often lack the ability to identify unknown AMR or other adaptive genes with novel mechanisms. To address this, we developed an integron gene cassette capture system, a functional screening platform that captures environmental gene cassettes for direct phenotypic testing. Using this system, we recovered previously unknown AMR determinants while also providing insights into the prevalence of known clinical AMR genes in a range of environmental samples, including food items. Here we provide experimental data on multiple novel bleomycin resistance genes and a stress response gene conferring gentamicin and tobramycin resistance. Our sequence analysis of the captured library also highlighted the diversity of the environmental cassette pool, with 656 unique cassettes recovered, the majority of which encoded proteins with unknown functions. The cassette capture system is a powerful tool for accessing hidden elements of the resistome and discovering novel adaptive genes that may go undetected using current sequence-based approaches.

**Environmental implication:** Antimicrobial resistance (AMR) genes are hazardous biological contaminants, yet the vast majority of environmental integron gene cassettes remain functionally uncharacterised. This study addresses this sequence-to-function gap by deploying a novel functional capture platform directly on realistic environmental matrices, including agricultural fertilisers, coastal seawater, and commercial food products. By characterising these cassettes, we uncovered hidden reservoirs of both novel and clinically established AMR genes circulating in critical exposure pathways. This work reveals the true hazardous potential of the mobile environmental resistome, validating a proactive One Health surveillance tool for monitoring emerging biological threats.

## 1. Introduction

The global emergence of antimicrobial resistance (AMR) is widely recognised as one of the most critical threats to public health worldwide, significantly undermining the efficacy of antibiotic treatments [1]. Increasingly, AMR is being understood not just as a clinical challenge, but as a significant issue of environmental pollution and One Health concerns across food production systems [2]. This crisis is accelerated by the rapid acquisition and spread of AMR genes via horizontal gene transfer (HGT) across diverse environments [3], with integrons playing a significant role in this process in clinically relevant Gram-negative pathogens [4]. Integrons are genetic elements capable of capturing and expressing exogenous mobile genes, known as gene cassettes, via a site-specific integrase (IntI) [5]. These cassettes typically consist of a promoterless open reading frame (ORF) and a recombination site (*attC*). IntI catalyses recombination between the *attC* site of a circularised gene cassette and the recombination site (*attI*) of the integron, allowing the accumulation of multiple cassettes into an array [6]. Expression of these captured genes is generally driven by a cassette promoter (Pc) located either within the *intI* coding sequence or between *intI* and *attI* site [4]. The activity of IntI is regulated by the bacterial SOS response, where cellular stress triggers expression of IntI, enabling cassette capture, rearrangement and excision. Integrons thus function as an aid to adaptive processes, providing bacteria with the ability to rapidly sample and acquire diverse adaptive functions, particularly during periods of stress [4,7,8].

Previous studies using phenotypic or metagenomic approaches show there is a vast diversity and abundance of integron gene cassettes in both human impacted and near-pristine ecosystems [9–12]. However, current analysis techniques provide limited information regarding the functional potential of the cassette metagenome, with ∼90% of environmental gene cassettes being typically of unknown function, largely due to a lack of sequence homology with genes of known function [10,11,13–15]. This sequence-to-function gap significantly limits our understanding of the environmental resistome, as our reliance on sequence similarity fails to detect novel resistance mechanisms [16].

To address this knowledge gap, we developed a novel assay for targeted capture and functional screening of environmental integron gene cassettes. This system exploits the structurally conserved site-specific recombination machinery of integrons to capture mobile gene cassettes from complex metagenomic DNA and place them under the control of an inducible promoter for direct phenotypic assessment. In this study, we recovered known clinical resistance genes, validating the system’s utility for AMR surveillance, while also identifying novel resistance determinants. The findings provide valuable insights into the undiscovered environmental resistome and demonstrate that functional metagenomic screening is essential for bridging the sequence-to-function gap to uncover novel threats to public health.

## 2. Methods

### 2.1. Bacterial strains, plasmids and culture conditions

The complete list of the bacterial strains and plasmids used in this study is provided in Table S1. *E. coli* strains were grown in Lysogeny Broth (LB) (Lennox) or on LB agar (LBA) supplemented with antibiotics where necessary. Mueller-Hinton (MH) broth was used for antimicrobial susceptibility testing. The final antibiotic concentrations used for selection were: 50 μg/mL kanamycin (Km), 100 μg/mL carbenicillin (Cb), and 10 μg/mL gentamicin (Gm). To support the growth of the auxotrophic *E. coli* strain WM3064 λpir [17], media were supplemented with 0.3 mM 2,6-diaminopimelic acid (DAP). Media were supplemented with sucrose to a final concentration of 5% (w/v) to select against *sacB*-containing plasmids. Induction of P*_ara_*, P*_cumate_* and P*_tac_* promoters was achieved by the addition of 2 mg/mL L-arabinose (Ara), 0.1 mM 4-Isopropylbenzoic acid (cumate), and 0.4 mM isopropyl β-D-1-thiogalactopyranoside (IPTG), respectively.

### 2.2. Construction of gene cassette capture system (pVR106-pUS250-*ccdB-attI1-ccdA* and pVR110-pttQ-*intI1*)

The gene cassette capture system was developed using a two-plasmid approach. Plasmid pVR106 was constructed as the capture vector, while pVR111 was designed to express IntI1 integrase. This system leverages the class 1 integron integrase (*intI1*) and its recombination site (*attI1*), coupled with a counterselection strategy to ensure the recovery of captured cassettes. We evaluated two counter-selection markers, *ccdB* toxin gene and *sacB* (conferring sucrose sensitivity) [18,19]. A series of in-frame *attI1* insertion variants were synthesised (Twist Bioscience, USA) to identify integration sites that preserved marker lethality (Table S2). These segments were cloned into the *Eco*RI-*Hin*dIII sites of pBAD24 [20] under the control of the arabinose-inducible promoter P_ara_. Except for one construct (*ccdB::attI1-5)*, all constructs retained full lethality. The *ccdB-attI1* variant pBAD24-*ccdB-attI1*-4, which harbours the *attI1* site at nucleotides 73–74 (codons I24–I25) of *ccdB*, was used for all subsequent experiments (Table S2). The PCR amplified 2,039-bp module (primers arapBAD-fow-*Bgl*II/arapBAD-rev-*Sal*I), comprising the P*ara* promoter and the *ccdB-attI1* construct, was assembled into the *Bgl*II- and *Sal*I-linearised pUS250 vector (Table S1) using NEBuilder HiFi DNA Assembly (NEB). To permit plasmid maintenance in the absence of induction, synthesised *ccdA* antitoxin gene was cloned into *Cla*I+*Sal*I sites under the control of the cumate-inducible promoter, generating the capture plasmid pVR106. The integron integrase source pVR110 (pttQ18-*intI1*), was constructed by assembling *intI1* (from plasmid R388) [21] into pttQ18 [22] using NEBuilder HiFi DNA Assembly (NEB) under the control of the IPTG-inducible P*_tac_* promoter. Plasmids pVR106 and pVR110 were transformed into *E. coli* MG1655, which served as the host strain for all subsequent assays.

To validate the capture system’s performance, we used metagenomic DNA, extracted from horse manure fertiliser, and prepared three types of DNA templates as input for the capture system: (i) total metagenomic DNA, (ii) PCR amplicons using primers HS286-HS287 to amplify gene cassettes [9], and (iii) the same amplicons treated with kinase and ligase to facilitate circularisation. After transformation and selection, we confirmed the captured cassettes via a combination of Sanger and Nanopore sequencing and identified 166 unique cassette arrays. Direct transformation of total metagenomic DNA yielded only a single gene cassette, and the linear amplicon approach yielded five. In contrast, the circularised amplicon strategy yielded 160 cassette arrays, that is consistent with the IntI1-mediated non-replicative recombination mechanism [23,24]. Given its high capture efficiency, the circularised amplicon strategy was adopted for all subsequent library construction in this study.

A technical constraint of this system was the emergence of escape mutants arising from the *ccdB* based counter-selection, a known challenge in toxin-based counter-selection where spontaneous mutations can readily inactivate the toxin under strong selective pressure [25–27]. We mitigated the background of non-recombinant colonies by directly subjecting the entire captured library to functional screening. Additionally, we employed automated colony picking using a PIXL microbial colny picker (Singer Instruments) for an efficient screening process. Source colonies were picked at 19 mm/s (9 mm PickupLine) and inoculated into LB+Km at 60 mm/s, paired with automated mixing (3 cycles of 3 revolutions). This pipeline effectively enriched for positive clones carrying the phenotypic traits of interest.

### 2.3. Construction of the synthetic gene cassette pVR111 (pUS321gm-*attC_aadA7_*_-bs_)

The donor plasmid for conjugation assays was constructed by annealing overlapping primers (*attC-aadA7*-fow/rev) designed to generate the *attC_aadA7_* sequence with *Nhe*I and *Xho*I overhangs (Table S3). This fragment was ligated into the mobilisable suicide vector pUS321 containing the constitutively expressed fluorescence marker *fuGFP* [28]. Additionally, the chloramphenicol resistance gene (*cat*) was replaced with the gentamicin resistance gene (*aacC1*) at the *Nde*I/*Kpn*I sites.

### 2.4. Construction of the expression control pVR106-*fuGFP*

To generate a positive control for cassette expression and a negative control for antibiotic screening, we synthesised the open reading frame of *fuGFP* (Twist Bioscience, USA) and assembled it into pVR106 using NEBuilder HiFi DNA Assembly (NEB). The *fuGFP* gene was positioned within the *attI1* recombination site to act as a captured promoterless cassette, conferring fluorescence upon induction.

### 2.5. Conjugation based *attC* × *attI1* recombination assays

Recombination efficiency between the *attC* site and the class 1 *attI1* sites was quantified using a suicide conjugation assay [29]. The donor strain in *E. coli* WM3064 λpir carrying synthetic gene cassette pUS321gm-*attC_aadA7_*_-bs_ (pVR111) was conjugated with recipient *E. coli* MG1655 [30] carrying pVR106 and pVR110 via filter mating in LBA+DAP. Expression of *intI1* was induced with 0.4 mM IPTG. Following a 6-hour incubation at 37°C, cells were recovered and plated on DAP-free LBA+Gm, LBA+Km and LBA+Km+Ara. Recombination frequencies were calculated as the ratio of recombinant colony forming units (CFUs) for Gm-resistant recombinants to total recipient CFUs from LBA+Km, or ratio of CFU for fluorescent colonies from LBA+Km+Ara to total recipient CFUs from LBA+Km after 2 days of incubation. Assays were performed in biological triplicates. Cointegrates were verified by colony PCR using primers *ccdB*-in-fow/rev (Table S3), followed by Sanger sequencing (Macrogen, South Korea).

### 2.6. Metagenomic DNA extraction and cassette amplification

To capture a broad diversity of gene cassettes, a set of environmental samples comprising bagged mixed salad leaves, Australian large green prawn digestive tracts, bagged horse manure fertiliser, and coastal seawater were collected (Table S4). Total metagenomic DNA extraction was performed using a FastPrep-24 (MP Biomedicals, USA) bead-beating protocol, as described previously [31,32]. To generate the DNA input for capture assays, integron gene cassette arrays were amplified from metagenomic DNA using primers HS286 and HS287 [9]. Additionally, class 1 integron cassettes were targeted using primer pairs HS915-HS459 and HS915-MRG285 [33,34]. Amplicons were subjected to phosphorylation treatment using T4 Polynucleotide Kinase (NEB) and blunt ligation using T4 DNA Ligase (NEB). Amplicons were purified using JetSeq™ Clean magnetic beads (Bioline).

### 2.7. Capture of gene cassette and functional screening for antimicrobial resistance genes

Purified cassette amplicons (50–200 ng) were electroporated into *E. coli* MG1655 cells harbouring pVR106 and pVR110. Transformants were recovered in 1 mL LB supplemented with 0.4 mM IPTG (to induce IntI1) and 0.1 mM cumate (to repress *ccdB*) for 2 hours at 37°C with shaking.

For functional screening, recovered library pools were incubated for 2 hours with 2 mg/mL L-arabinose to induce expression of captured cassettes. Cultures were plated onto LBA+Ara with the antibiotic of interest at concentrations that are inhibitory to the control strain *E. coli* MG1655 (pVR106-*fuGFP*). Details of the antimicrobials and their concentrations are provided in Table S5. Resistant colonies were screened via colony PCR (primers *ccdB*-in-fow/rev) to confirm cassette insertion (Table S3). PCR validated clones were sequenced via Sanger or Oxford Nanopore sequencing and re-transformed into fresh *E. coli* MG1655 to further phenotypically confirm the resistance conferred by each gene.

### 2.8. Quantitative minimal inhibitory concentration determination

Minimum inhibitory concentrations (MICs) were determined for each clone using the broth microdilution method [35]. Overnight cultures were subcultured (1:10 dilution) and induced with 2 mg/mL L-arabinose for 2 hours at 37°C. Cultures were then adjusted to a density of 5×10^5^ CFU/mL and inoculated into 96-well microtiter plates containing MH+Km+Ara broth further supplemented with the test antibiotic at varying concentrations. Plates were sealed with breathable films and incubated at 37℃ for 24 hours in a PHERAstar FSX plate reader (BMG Labtech). All experiments were performed using at least three biological replicates.

### 2.9. Nanopore sequencing of cassette amplicon pools and DNA sequence analysis

Plasmid DNA from the cassette library pool was extracted using the Monarch Plasmid Miniprep Kit (NEB). The library was PCR amplified using the junction primers *ccdB*-in-fow/rev. Amplicons were sequenced on a MinION Mk1B device using an R10.4.1 flow cell and Ligation Sequencing Kit (SQK-LSK114). Basecalling and demultiplexing were performed using Dorado v1.0.1 (https://github.com/nanoporetech/dorado) with the super high-accuracy model.

Sequence processing was performed to identify reads representing the plasmid backbone with a valid cassette inserted at the *attI1* site. For this, reads were first trimmed and oriented based on the forward and reverse *ccdB* junction primers using Pychopper v2.5.0 (https://github.com/epi2me-labs/pychopper) [parameters: -m edlib], and filtered for a minimum read length of 250 bp and a mean quality score of Q15 using Chopper v0.10.0 [36] [parameters: -q 15 -l 250]. Filtered reads were corrected using Canu v2.1.1 [37] [parameters: -correct -nanopore genomeSize=5000 corMhapSensitivity=high stopOnLowCoverage=1 corMinCoverage=0 corOutCoverage=100000 maxInputCoverage=100000 minInputCoverage=1 minReadLength=250 minOverlapLength=200 useGrid=false]. Corrected reads were then screened for the 50bp of sequence flanking each side of the *attI* insertion point using BLASTn v2.15.0 [38] [parameters: -task blastn -word_size 7 -dust no]. BLASTn results were filtered for a minimum alignment length of 45 bp and 90% sequence identity, specifically retaining only reads that had exactly one copy of each left-hand and right-hand flanking sequence. Reads were then clustered at 95% nucleotide identity with Vclust v1.2.3[39] using the graph-based Leiden clustering algorithm [40] [parameters: --algorithm leiden --metric tani --tani 0.95 --qcov 0.9 --rcov 0.9]. A single consensus sequence was generated for each cluster using SPOA v4.1.4 [41] [parameters: -r 0], and polished using Medaka v2.1.0 (https://github.com/nanoporetech/medaka) [parameters: medaka_consensus --bacteria]. From each consensus sequence, the specific region representing the inserted cassette was extracted using BEDTools v2.31.0 [42]. Open reading frames (ORFs) were predicted from the extracted regions using Prodigal v2.6.3 [parameters: -p meta] [43].

### 2.10. Functional annotation and structural bioinformatics

Functional annotation of the predicted protein sequences was conducted using eggNOG-mapper v2.0.1b (DIAMOND mode) [44–46], InterProScan v5.59-91.0 [47,48] and NCBI conserved domain search using Batch CD-Search [49], all using default parameters. Additionally, to account for any potentially unpredicted ORFs missed by Prodigal, we performed translated sequence searching of the complete inserted nucleotide sequences against the UniRef50 database[50] using MMseqs2 [51] with default parameters. Putative antibiotic resistance gene cassettes were also predicted using AMRFinderPlus [52] and RGI v6.0.6 against the CARD database [53], and by searching against the ResFinder database [54] using MMseqs2 with default parameters. Cassettes encoding transmembrane or secreted proteins were identified by screening for signal peptides with SignalP 6.0 [55] [parameters: -org arch|gram+|gram--format short]. IntegronFinder 2.0.5 [56] was used to identify *attC* sequences [parameters: --local-max --func-annot –gbk].

Three-dimensional protein structures were predicted using AlphaFold3 [57], with model confidence assessed via the interface Predicted Template Modelling (ipTM) score. The predicted models were queried against the Protein Data Bank (PDB) [58] and UniProt-AlphaFoldDB [59–61] using the Foldseek server [62] to identify structural homologs. Structural superimposition figures were generated using the Foldseek web interface.

## 3. Results

### 3.1. Development of an integron cassette capture system

We developed an integron cassette capture system for the recovery and direct functional screening of environmental gene cassettes (Figure 1A). The core principle of the system relies on a counter-selection strategy utilising the *ccdB* lethal toxin gene. We incorporated the recombination site (*attI1*) directly into the *ccdB* coding sequence without disrupting toxin function, placing this chimeric *ccdB::attI1* construct under the control of an L-arabinose inducible promoter (Para) (Methods). This design creates a survival dependency. When a diverse pool of circularised gene cassette amplicons (e.g., derived from environmental DNA) is introduced, IntI1-mediated recombination integrates a cassette into the *attI1* site. This integration physically disrupts the *ccdB* open reading frame. Consequently, upon L-arabinose induction, cells lacking a captured cassette perish due to toxin production, while those harbouring a successful integration survive.

**Figure 1.**
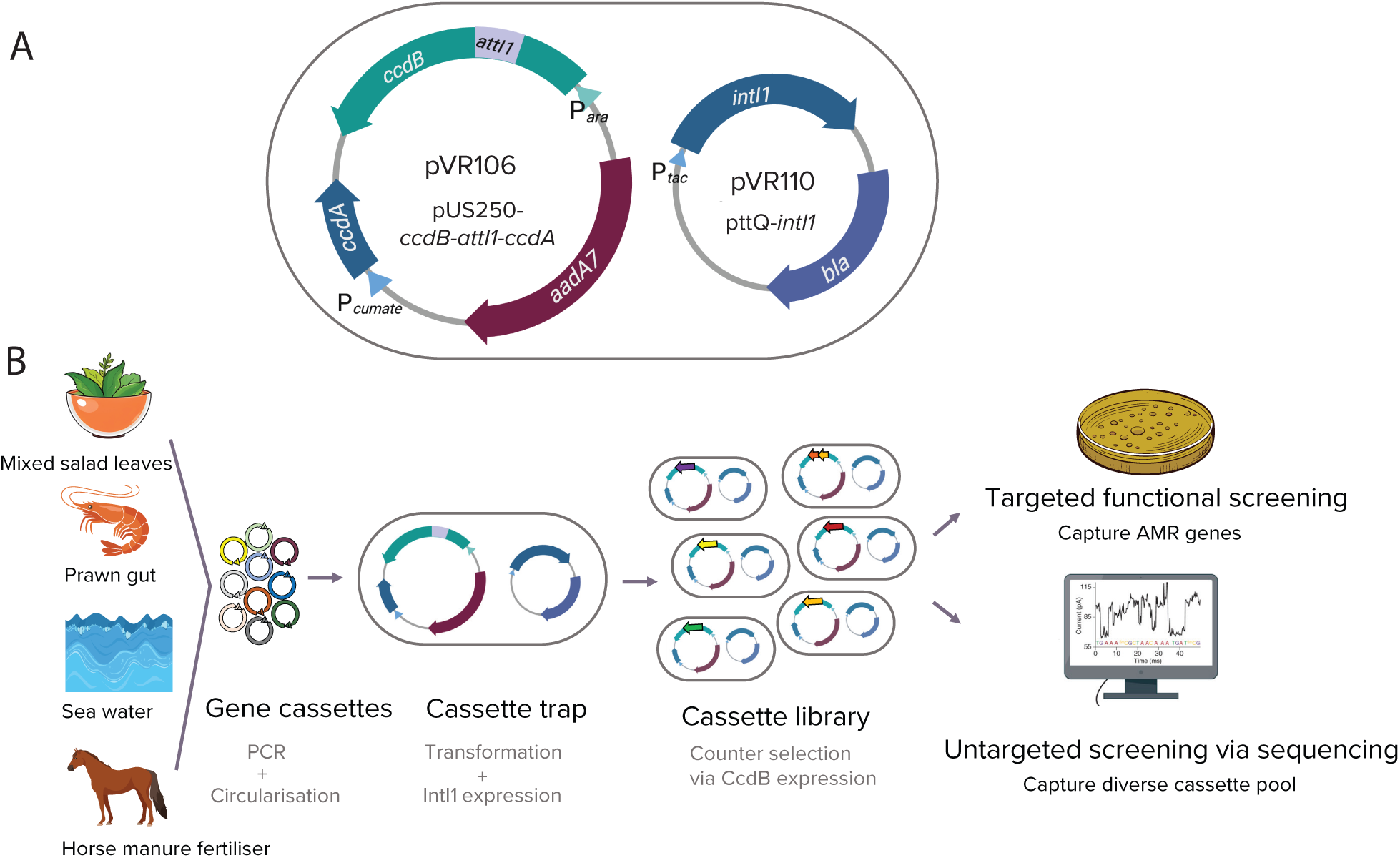
Schematic of the integron gene cassette capture system and experimental workflow. (A) Plasmid maps of the cassette capture system. The capture vector, pVR106 (pUS250*-ccdB-attI1-ccdA*) (Table S1), contains a *ccdB* gene, with an *attI1* recombination site embedded within it, under the control of an L-arabinose inducible promoter, allowing counter-selection of successful, site-specific insertion events. Additionally, this plasmid contains the antitoxin gene *ccdA* under a cumate-inducible promoter to protects cells against the toxic CcdB. Plasmid pVR110 (pttQ-*intI1*) contains *intI1* under the control of an IPTG-inducible promoter. (B) Workflow for the recovery and characterisation of the cassette metagenome from different environments. Gene cassettes were amplified from diverse environmental DNA sources (mixed salad leaves, prawn gut, sea water, and horse manure fertiliser), circularised and transformed into recipient cells expressing IntI1. Integration of a cassette into the *attI1* site disrupts the *ccdB* reading frame, ensuring that cells containing a captured cassette survive counter-selection. The resulting library was subjected to two parallel analyses: targeted functional screening to identify AMR genes, and untargeted nanopore sequencing of the PCR-amplified plasmid DNA (using *ccdB*-in-fow/rev junction primers) to characterise the diversity of the captured cassette pool. Schematic created with BioRender.com and modified using assets from Adobe Stock.

By simultaneously eliminating empty vectors, this strategy generates a complex library of transformed cells, each harbouring a gene cassette. Furthermore, because the captured cassettes are integrated directly downstream of the P*ara* promoter, the surviving library is immediately ready for functional expression screening. This approach bypasses the need for individual gene sub-cloning and provides a more targeted method for sampling the mobile gene pool than traditional approaches relying on large insert, fragmented metagenomic libraries.

To quantify the recombination efficiency of the capture system, we performed a suicide conjugation assay [63]. As a donor, we used the circularised gene cassette pVR111 (pUS321gm-*attC*aadA7-bs) encoding a gentamicin resistance determinant and a fluorescence marker (Table S1). This assay yielded a recombination frequency of 1.54 × 10⁻² when selecting for the gentamicin resistance marker, and 1.79 × 10⁻² when selecting on L-arabinose induced media. These efficiencies are consistent with previously reported values for IntI1-mediated recombination between *attI1-attC_aadA7_* [29]. We then validated the system’s performance in its intended direct electroporation workflow (Figure 2B). When transformants were selected against the cassette encoded gentamicin resistance marker, 99% of the resulting colonies contained a successful insertion while selection based on inducing *ccdB* activity yielded a 40% success rate, and the remaining background colonies consisted of *ccdB* escape mutants or had insertions of host chromosomal IS element *IS*2. The issue of high escape mutant rate was bypassed by subjecting the entire captured library to direct functional screening, and by using automated colony handling to increase throughput (Methods).

**Figure 2.**
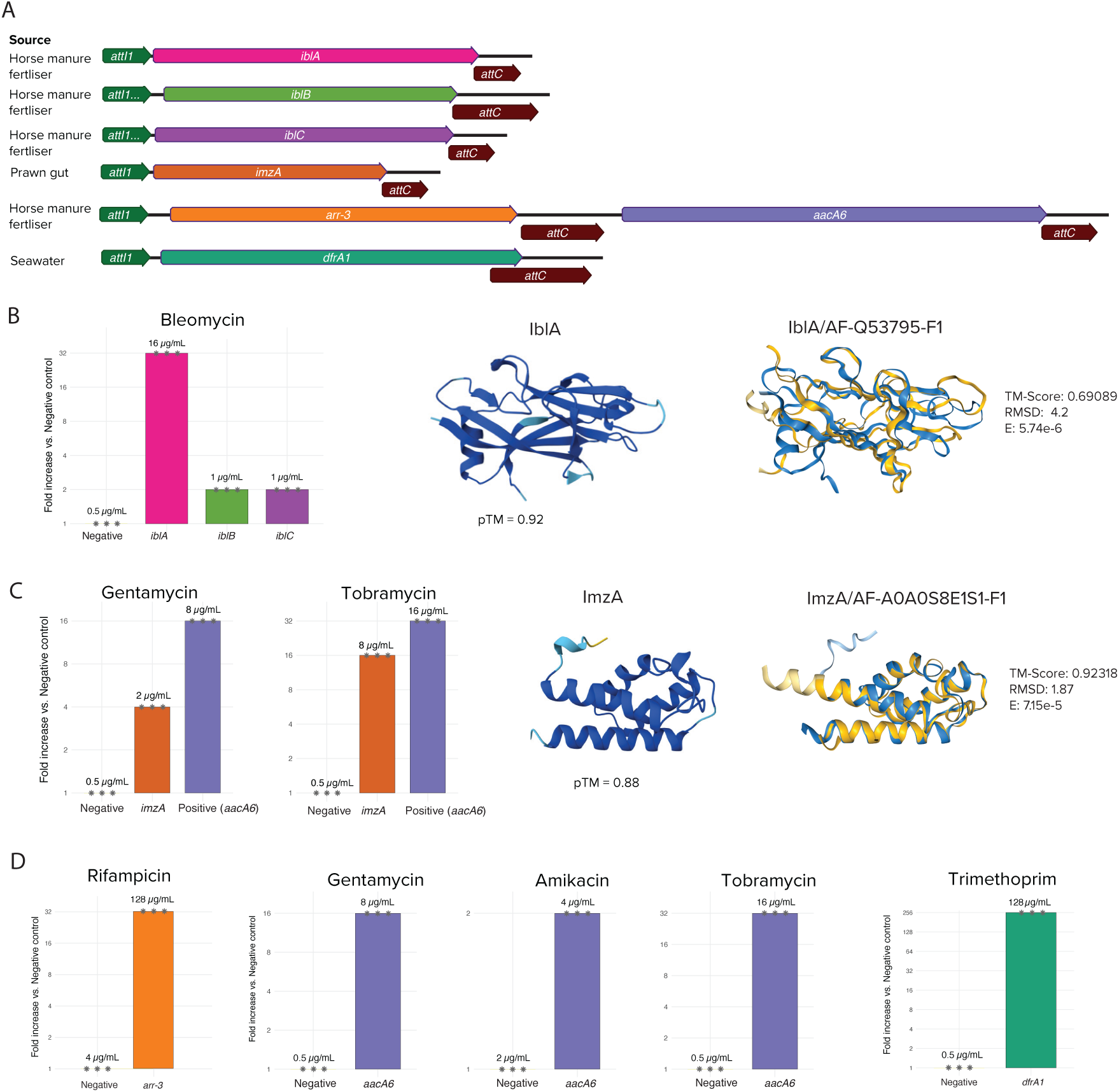
Functional validation of antimicrobial resistance gene cassettes recovered from environmental samples. (A) Captured gene cassette arrays and their corresponding environmental sources. Long arrows indicate open reading frames (ORFs) and their transcriptional orientation, maroon arrows represent *attC* sites and short green arrows represent the *attI1* site. (B) Antimicrobial resistance (AMR) profile of bleomycin resistance genes, AlphaFold3 model of IblA and structural superimposition of IblA (blue) onto its structural homolog AF-Q53795-F1 (yellow). (C) AMR profile of ImzA, AlphaFold3 model and structural superimposition of ImzA (blue) onto its structural homolog AF-A0A0S8E1S1-F1 (yellow). (D) AMR profiles for the known AMR-conferring genes (in parentheses): rifampicin (*arr-3*), aminoglycoside acetyltransferase (*aacA6*), and trimethoprim (*dfrA1*). Bar charts display the fold-increase in minimum inhibitory concentration (MIC) relative to the negative control (pVR106-*fuGFP*) (Table S1). The antimicrobial concentrations (µg/mL) are indicated above each bar, and biological replicates (n = 3) are represented by asterisks.

### 3.2. Functional metagenomic screening of captured cassette libraries identified novel antibiotic resistance genes

We investigated the utility of our capture system, coupled with functional selection, in identifying gene cassettes conferring antibiotic resistance across different samples. For this, gene cassette libraries were generated using circularised amplicons of DNA from environmental sources: supermarket produce (mixed salad leaves, fresh prawn gut), horse manure fertiliser and coastal seawater (Table S4). For each sample, a library containing captured cassettes was generated and was subjected to functional selection against a panel of antimicrobials (Table S5). This approach resulted in the subset of plasmids carrying inserted gene cassettes capable of conferring resistance to each tested antibiotic being retained while eliminating captured cassettes that did not confer AMR phenotype of interest (Figure 1B).

The functional screen identified three novel gene cassettes that conferred resistance to bleomycin, designated *iblA* (integron-derived bleomycin resistance A), *iblB* and *iblC*, captured from horse manure fertiliser sample (Figure 2). Quantitative MIC assays revealed that *iblA* conferred a 32-fold increase in resistance (MIC 16 µg/mL), while *iblB* and *iblC* conferred a 2-fold increase (MIC 1 µg/mL) relative to the negative control (MIC 0.5 µg/mL). The *iblA* determinant was identified as a 426 bp ORF within a 483 bp gene cassette. IblA shares 89.9% amino acid identity with a hypothetical protein from a previously reported environmental gene cassette [9], but lacks homology to any characterised resistance protein family. Conserved domain searches identified only a domain of unknown function DUF5990 (Pfam19452). Structural homology searches using AlphaFold3/Foldseek [57,62] revealed a structural alignment with a bleomycin/phleomycin-binding protein and ankyrin-like bleomycin resistance protein from *Streptomyces verticillus* (TM-score = 0.69089, RMSD = 4.2, E-value = 5.74e-6) (Figure 2) [64]. The *iblB* (384 bp ORF in 507 bp cassette) and *iblC* (390 bp ORF in 450 bp cassette) encoding proteins were identified as new members of the Vicinal Oxygen Chelate (VOC) superfamily (COG3324). IblB shared 78.74% amino acid identity with a VOC family protein from *Pseudomarimonas* sp. (GenBank GCA_052390415), while IblC shared 63.71% amino acid identity with a VOC homolog from *Burkholderia alba* (GenBank GCF_034721475). Functional annotation using the EggNOG database [44,45] classified both proteins as glyoxalase/bleomycin resistance proteins (Table S6).

Screening for aminoglycoside resistance identified another novel determinant, designated *imzA* (integron derived MazG-like protein A), captured from a prawn gut sample (Figure 2). This cassette conferred a 4-fold increase in the MIC for gentamicin (MIC 8 µg/mL) and a 16-fold increase for tobramycin (MIC 8 µg/mL), relative to the control (MIC 2 µg/mL and 0.5 µg/mL, respectively). *imzA* was identified as a 306 bp ORF within a 365 bp gene cassette. ImzA shared no sequence homology with any known aminoglycoside modifying enzymes or 16S rRNA methyltransferases. We found a similar hypothetical protein with 81% identity detected in *Nitrosomonas eutropha* (GenBank GCA_900116685), which itself lacks functional annotation. Structural homology searches using AlphaFold3/Foldseek identified a high confidence structural alignment (TM-score = 0.923, RMSD = 1.87, E-value = 7.15e-5) with a protein from a *Planctomycetes* spp. annotated as a putative NTP pyrophosphohydrolase MazG (AF-A0A0S8E1S1-F1) (Figure 2). Past work indicates that MazG family NTP pyrophosphohydrolases are involved in regulating bacterial stress adaptation responses [65]. Subcellular localisation predictors returned no signal peptides or transmembrane domains and predicted a cytoplasmic localisation, consistent with a soluble cytoplasmic protein.

In addition to these novel genes, functional screening recovered well characterised, class 1 and class 2 integron-associated antibiotic resistance cassettes. For example, screening of the horse manure fertiliser sample identified a 1,241 bp insert containing two adjacent gene cassettes in a typical cassette array arrangement (Figure 2). This array conferred a >32-fold increase in resistance to rifampicin (>128 µg/mL), a 32-fold increase to tobramycin (16 µg/mL), a 16-fold increase to gentamicin (8 µg/mL), and a 2-fold increase to amikacin (4 µg/mL) relative to the negative control group. Sequence analysis revealed 100% nucleotide identity to cassette arrays found on class 1 integrons from various *Acinetobacter* spp. isolated from clinical settings (e.g. *A. baumannii*, *A. lwoffii*; GenBank CP078043, KU133345). Of the two genes, the first shared 99.9% amino acid identity with a rifampicin ADP-ribosyltransferase (*arr-3*) from *Klebsiella pneumoniae* (GenBank AGO62693), and the second shared 99.48% identity with an aminoglycoside N(6’)-acetyltransferase (*aacA6*) reported in *Enterobacter cloacae* CY01 (GenBank AIK02012) (Figure 2).

Additionally, we captured a 578 bp gene cassette from a seawater sample conferring resistance to trimethoprim with a >256-fold increase in MIC (>128 µg/mL) relative to the control (0.5 µg/mL) (Figure 2). This cassette contained the dihydrofolate reductase gene *dfrA1*, sharing 99.8% amino acid identity to DfrA1 found in class 2 integrons (GenBank JX867127). *dfrA1* is widely disseminated across numerous Gram-negative pathogens primarily isolated from clinical samples, including *Proteus vulgaris* (GenBank JX867127), *Proteus mirabilis* (GenBank CP045257, KY885013), *Klebsiella pneumoniae* (GenBank CP169615, CP061064), *Citrobacter freundii* (GenBank CP054294.1), *Escherichia coli* (GenBank OZ038647), and *Vibrio cholerae* (GenBank CP189152) frequently as part of class 2 integrons [66].

### 3.3. Captured cassette libraries from all environments hosted diverse suites of novel gene cassettes

After demonstrating that functional screening enriches a pool of gene cassettes that encode resistance to diverse antimicrobials, we explored the broader functional diversity captured by the system. Sequencing of the entire mixed pools of captured cassettes from each sampled environment was followed by bioinformatic analysis using pipelines adapted from our previous work [13] to identify reads containing genuine gene cassettes (reads showing a valid cassette insertion at the *attI1* site in the capture plasmid backbone with at least one *attC* site). The analysis yielded a total of 656 gene cassettes from the four input DNA sets (Figure 3).

**Figure 3.**
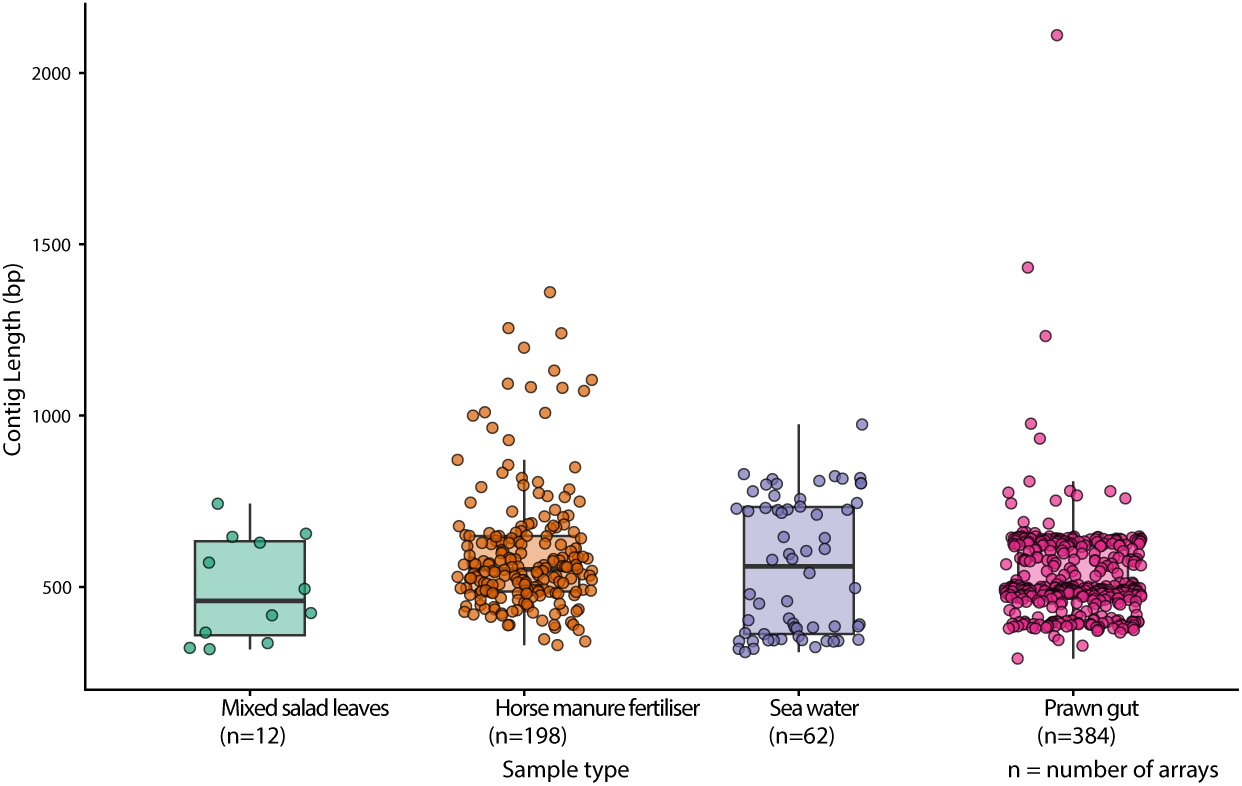
Distribution of captured gene cassette array lengths across different environmental sources. Box plots displaying the size distribution (in bp) of unique gene cassette arrays recovered from mixed salad leaves (n=12), horse manure fertiliser (n=198), seawater (n=62), and prawn gut (n=384) cassette metagenomes. Each point represents a distinct cassette array assembled from Nanopore sequencing reads.

The set of captured gene cassettes were first subject to functional prediction using NCBI Conserved Domain search, InterProScan[47,48], EggNOG-mapper [44,45], UniRef50 database [50], ResFinder database [54], AMRFinderPlus [52], RGI against the CARD database [53] and DefenseFinder [67]. This analysis assigned putative functions to 8% of the captured gene cassettes (Table S6). The remaining were either not assigned a function or were classified as hypothetical proteins. This finding is consistent with previous investigations into gene cassette-encoded functions [10,11,13–15,68]. While only a small proportion were functionally annotated, this set did contain many functions of interest, including genes bioinformatically predicted to play a role in microbial adaptation, antibiotic resistance, toxin-antitoxin systems, anti-phage defence, virulence and stress tolerance functions (Table S6).

The untargeted capture approach again identified genes conferring resistance to trimethoprim (*dfrA1*), aminoglycosides (*aacA6*), rifampicin (*arr-3*) as well as three genes encoding members of the glyoxalase/bleomycin resistance family which belong to the vicinal oxygen chelate (VOC) superfamily (COG3324) (Table S6). This included a subset of the AMR genes recovered during the targeted functional screening, including novel VOC family genes (*iblB* and *iblC*), as well as additional genes of interest. Untargeted screening using DNA from horse manure fertiliser samples revealed two cassettes encoding proteins with a P-loop NTPase domain (IPR027417), a domain noteworthy as it was recently characterised in Gar, a novel aminoglycoside modifying enzyme identified through a similar functional metagenomic screening approach [16]. Although the two proteins identified in this study share no sequence similarity with *gar*, their shared domain architecture suggests potential for a similar function. This sample also contained two cassettes encoding a monooxygenase domain (Pfam03992), which is reported to be involved in antibiotic production [69].

Toxin-antitoxin (TA) systems are commonly reported in integron gene cassettes, and we identified a cassette encoding the ParE toxin of a type II TA system (pfam05016) and a type II antitoxin from the HicB family from horse manure fertiliser and seawater samples respectively (Table S6). These TA systems have previously been found associated with gene cassettes from *Vibrio* spp. and human associated bacterial communities [68,70,71]. Additionally, we found integron cassette genes potentially involved in defence against bacteriophages. In the horse manure fertiliser sample, DefenseFinder identified a gene cassette with significant homology to gcuWGS21 (E-value = 2.0e-19; score = 55.9), a recently described anti-phage defence system found within gene cassettes [72]. In another case, a cassette recovered from the seawater sample encoded a gene containing a HEPN domain (Pfam18739), frequently associated with RNase activity in phage defence systems [73]. We also identified a gene encoding a Type IV secretion system component (VirB/VirD4) from the seawater sample, which has previously been reported in gene cassettes [74,75].

We also identified cassettes carrying genes that are implicated in tolerance to abiotic stressors and detoxification of chemical compounds (Table S6). This includes four cassettes from horse manure fertiliser samples that encode glutathione dependent formaldehyde activating enzymes (Pfam04828). Genes with this putative function have previously been observed in gene cassettes from natural environments in metagenomic analyses [68]. These enzymes are involved in detoxifying formaldehyde, a common metabolic byproduct and environmental toxin [76,77]. Additionally, this sample was found to contain three non-identical cassettes encoding the quinol monooxygenase YgiN implicated in mitigating reactive oxygen species [78]. While a cassette containing a CspA family cold shock protein (Pfam00313) potentially involved in adaptation to low temperatures [79] was recovered from the seawater sample.

## 4. Discussion

Integrons play a key role in facilitating the acquisition and dissemination of accessory genes, including those conferring AMR [80]. While sequence-based surveys have revealed a vast and diverse pool of environmental gene cassettes [10,81], the inability to functionally characterise their encoded products has severely limited our understanding of the environmental resistome.

By deploying our cassette capture system, we identified multiple novel AMR genes from diverse environmental sources. Two of these bleomycin resistance genes, *iblB* and *iblC* encode protein products belonging to the Vicinal Oxygen Chelate (VOC) superfamily, a group we recently confirmed is enriched within integron gene cassette libraries [11]. A third bleomycin resistance gene *iblA* was even more divergent, sharing no significant sequence homology with known resistance proteins. Its function could only be inferred through protein structural analysis which revealed alignment with a bleomycin-binding protein [64]. For each of these, activity was demonstrated and MICs determined. Bleomycin is a glycopeptide antibiotic used in chemotherapy [82] and the detection of multiple, novel resistance determinants underscores the role of the environmental cassette metagenome as a reservoir for clinically relevant traits.

We also identified an additional aminoglycoside resistance gene, *imzA*. While the level of resistance to aminoglycosides gentamicin and tobramycin conferred by this gene was lower than for the canonical aminoglycoside modifying enzyme encoded by *aacA6*, the mode of action appears to be novel. ImzA showed no sequence or predicted structural similarity to any currently characterised aminoglycoside modifying enzymes, 16S rRNA methyltransferase or other characterised resistance determinant. Structural homology analysis identified high-confidence similarity between ImzA and a MazG family NTP pyrophosphohydrolases (UniProt P96379), which regulate bacterial stress adaptation responses [65,83,84]. This structural conservation suggests ImzA may function in stress response regulation rather than direct aminoglycoside modification. Aminoglycosides are known to trigger broad cellular stress responses in bacteria [85], and the observed resistance phenotype may reflect a stress response-mediated tolerance. Further biochemical studies, including assessment of stress response activation during aminoglycoside exposure will help elucidate the precise molecular mechanism of ImzA.

In addition to capturing novel gene, this system also recovered widely disseminated, clinically important AMR determinants. The presence of these genes in our environmental samples highlights the overlapping resistomes of clinical and non-clinical settings and underscores the importance of One Health focussed research. Successfully capturing these known markers provides a vital proof of concept for the system’s role in monitoring antibiotic resistance as an environmental pollutant.

While techniques for identifying integron gene cassettes in metagenomes [10,11,13] and diverse environmental hosts [86–88] continue to expand, gene cassette functional characterisation has remained extremely limited due to sequence novelty and the challenges of experimental validation. Our findings highlight the power of our combined cassette capture and functional screening tool to begin to address this, particularly in identifying AMR genes which utilise novel mechanisms. Detecting such divergent AMR determinants is important for clinical practice. This was recently highlighted by work using an alternative functional metagenomic approach which led to the characterisation of a novel aminoglycoside resistance gene present in clinical isolates that had remained undetected due to its sequence novelty [16]. Reliance on current sequence-based surveillance methods risks hidden reservoirs of resistance circulating undetected until they become clinically problematic. Our system is therefore timely, providing functional screening of environmental gene cassettes in a manner well suited for revealing novel resistance determinants that may otherwise be missed.

Beyond AMR, this approach can identify additional clinically important phenotypes disseminated by gene cassettes. Recent efforts to functionally characterise integron gene cassettes have led to the important discovery that numerous integron gene cassettes carry novel anti-phage defence genes [72,89,90]. Systematic cloning and screening of gene cassettes of unknown function revealed that integrons serve as a reservoir for novel anti-phage defence systems, a finding of clinical relevance given the growing interest in phage therapy [72]. Here, our approach identified one of these recently discovered phage defence system gcuWGS21 [72]. These findings emphasise the importance of methods for rapid identification of cassettes carrying adaptive and potentially clinically relevant genes, particularly those with novel mechanisms of action, prior to their widespread dissemination.

Our tool also has potential to extend our understanding of integron gene cassette functional roles. Sequencing of the entire mixed pool of captured cassettes provided broader functional insights. While most (∼92%) of captured cassettes encoded hypothetical proteins, consistent with other large-scale surveys of environmental integrons [10,11,15,68], functional predictions were made for 53 cassettes. In addition to toxin-antitoxin systems and anti-phage defence gene cassettes, functions now well associated with integrons, we identified genes cassettes with predicted roles in stress tolerance. This includes a cassette conferring adaptation to low temperatures and four cassettes encoding formaldehyde detoxification functions [68,91]. This is consistent with the view that integrons are general platforms for acquiring genes conferring diverse functions, extending beyond AMR determinants [11,13].

In conclusion, this work presents a new platform for the functional exploration of the integron gene cassette metagenome. Using this system, we show that the environmental gene cassette pool contains a variety of novel antibiotic resistance genes not detectable by sequence-based methods. Our findings reinforce the importance of integrons as platforms for mobilising diverse adaptive traits. The cassette capture system provides a powerful tool for functional screening, enabling the discovery of novel antimicrobial resistance genes and other novel biological functions from diverse environmental sources. Application of such approaches are urgently needed to better understand the environmental resistome and aid in anticipating future threats to public health.

## Author contributions

**Study conceptualisation:** VR, TMG, QQ, MRG, NC, SGT. **Experimental procedures:** VR, DHR, CS, VJM **Data analysis:** VR, TMG, EC, BS **Funding acquisition:** SGT. **Writing, review & editing:** All authors.

## Acknowledgements

Dr. Vaheesan Rajabal is supported by funding from the ARC Centre of Excellence in Synthetic Biology (CE200100029).

## Data availability

Data will be made available on request.

